# Targeting the translational machinery in gastrointestinal stromal tumors (GIST) – a new therapeutic vulnerability

**DOI:** 10.1101/2021.09.01.458633

**Authors:** Donna M. Lee, Angela Sun, Sneha S. Patil, Lijun Liu, Aparna Rao, Parker Trent, Areej A. Ali, Catherine Liu, Jessica L. Rausch, Laura Presutti, Adam Kaczorowski, Felix Schneider, Nduka Amankulor, Masahiro Shuda, Anette Duensing

**Author notes:** <u>Corresponding author:</u> Anette Duensing, UPMC Hillman Cancer Center, Research Pavilion, Suite G.17, 5117 Centre Avenue, Pittsburgh, PA 15213, USA. Co-first authors (Donna M. Lee, Angela Sun). Department of Neurological Surgery, The University of Pennsylvania, 3 Silverstein Pavilion (Nduka Amankulor). The authors declare that they have no competing financial interests.

## Abstract

Although *KIT*-mutant GISTs can be effectively treated with tyrosine kinase inhibitors (TKIs), many patients develop resistance to imatinib mesylate (IM) as well as the FDA-approved later-line agents sunitinib, regorafenib and ripretinib. Resistance mechanisms mainly involve secondary mutations in the *KIT* receptor tyrosine kinase gene indicating continued dependency on the KIT signaling pathway. The fact that the type of secondary mutation confers either sensitivity or resistance towards TKIs and the notion that secondary mutations exhibit intra- and intertumoral heterogeneity complicates the optimal choice of treatment in the imatinib-resistant setting. Therefore, new strategies that target KIT independently of its underlying mutations are urgently needed. Homoharringtonine (HHT) is a first-in-class inhibitor of protein biosynthesis and is FDA-approved for the treatment of chronic myeloid leukemia (CML) that is resistant to at least two TKIs. HHT has also shown activity in *KIT*-mutant mastocytosis models, which are intrinsically resistant to imatinib and most other TKIs. We hypothesized that HHT could be effective in GIST through downregulation of KIT expression and subsequent decrease of KIT activation and downstream signaling. Testing several GIST cell line models, HHT led to a significant reduction in nascent protein synthesis and was highly effective in the nanomolar range in IM-sensitive and IM-resistant GIST cell lines. HHT treatment resulted in a rapid and complete abolishment of KIT expression and activation, while KIT mRNA levels were minimally affected. The response to HHT involved induction of apoptosis as well as cell cycle arrest. The antitumor activity of HHT was confirmed in a GIST xenograft model. Taken together, inhibition of protein biosynthesis is a promising strategy to overcome TKI resistance in GIST.

## INTRODUCTION

The majority of gastrointestinal stromal tumors (GIST) arise from oncogenic mutations in the *KIT* receptor tyrosine kinase gene^1–3^. Hence, most inoperable and/or metastatic GISTs can be treated with imatinib mesylate (IM; Gleevec®), a small molecule tyrosine kinase inhibitor (TKI) that targets KIT and leads to disease stabilization in more than 85% of the patients^4,5^. However, complete remissions are rare, and approximately 50% of the patients experience disease progression within the first two years of therapy^6^. Drug resistance is mostly due to secondary *KIT* mutations that prevent imatinib from binding^7,8^. Unfortunately, second- and third-line treatments, the TKIs sunitinib (Sutent®) and regorafenib (Stivarga®), only offer limited additional benefit^9–11^. A fourth-line medication (ripretinib, Qinlock®) was recently approved, but development of resistance is still a problem^12^. Another small molecule inhibitor, avapritinib (Ayvakit®), was approved only for a subset of GIST that carry a certain *PDGFRA* (platelet-derived growth factor receptor alpha) mutation (*PDGFRA* D842V)^13,14^. Hence, new therapeutic approaches for GIST patients are still needed, in particular when TKI resistance has occurred.

GIST cells are highly dependent on KIT signaling for their survival and proliferation. This is exemplified by the fact that GIST cell apoptosis is not only induced by TKI-mediated inhibition^4,15^, but also by loss of KIT expression via other mechanisms, such as siRNA-mediated knockdown, treatment with HSP90 inhibitors or inhibition of *KIT* gene transcription^16–19^. In addition, resistance mechanisms mainly involve secondary *KIT* mutations that prevent drug binding, further pointing towards the continued dependency on an activated KIT signaling pathway^7,8^. Because secondary mutations exhibit heterogenous sensitivity to later-line TKIs and also display intra- and intertumoral heterogeneity, treatment responses are likely to be uneven depending on the clonal composition of the tumor^20^. Therapeutic strategies that target KIT independently of its underlying mutation could potentially overcome these obstacles^21^. In this scenario, targeting KIT protein expression by inhibiting mRNA translation represents a promising therapeutic strategy^22^.

Homoharringtonine (HHT) is a cytotoxic alkaloid that is derived from the Japanese Plum Yew (cephalotaxus harringtonia)^23–25^. It was originally used in Chinese traditional medicine and is now produced as semisynthetic, purified compound (omacetaxine mepesuccinate/Synribo®; Teva Pharmaceuticals)^22,26^. HHT acts as a global protein translation inhibitor by interfering with the elongation step of protein biosynthesis. It competitively binds the peptidyl transferase A-site cleft of ribosomes, thereby preventing the incoming amino acid side chains of aminoacyl-tRNAs from attaching^24,27^. HHT/omacetaxine is FDA-approved for the treatment of patients with chronic myeloid leukemia (CML) that are resistant and/or intolerant to at least two TKIs^26,28^. Further clinical and preclinical studies have shown HHT’s activity in acute myeloid leukemia, *VHL*-mutant clear cell renal cell carcinoma as well as rhabdoid tumors^25,29,30^. Of note, HHT has also shown preclinical activity in *KIT* D816V-mutant mastocytosis models, a disease that is intrinsically resistant to imatinib and most other TKIs targeting KIT^31^.

Our study investigates whether HHT can target KIT in GIST cells by inhibiting protein translation. We hypothesized that treatment with HHT reduces KIT protein levels, thereby decreasing KIT activation as well as its downstream signaling cascades ultimately leading to GIST cell apoptosis in an imatinib-sensitive as well as imatinib-resistant setting.

## MATERIALS AND METHODS

### Cell culture, inhibitor treatments

The imatinib-sensitive human GIST cell lines GIST882 (homozygous mutation in *KIT* exon 13 pK642E; kindly provided by Jonathan A. Fletcher, Brigham and Women’s Hospital, Boston, MA, USA) and GIST-T1 (heterozygous *KIT* exon 11 deletion pV560_Y578del; kindly provided by Takahiro Taniguchi, Kochi Medical School, Kochi, Japan) were derived from untreated metastatic GISTs^32,33^. Imatinib-resistant GIST cell lines GIST430 (heterozygous primary *KIT* exon 11 deletion [pV560_L576del], secondary *KIT* exon 13 point mutation [pV654A]), GIST48 and GIST48B (both: homozygous primary *KIT* exon 11 point mutation [pV560D], secondary *KIT* exon 17 point mutation [pD820A]; all kindly provided by Jonathan A. Fletcher, Brigham and Women’s Hospital, Boston, MA, USA) were derived from human GISTs that had developed clinical resistance to imatinib therapy^18,19^. GIST48B is a spontaneous subline derived from GIST48 (same *KIT* mutational status) but does not express KIT mRNA and protein at appreciable levels^34^. All GIST cells were propagated as previously described^15,18,19^.

Homoharringtonine (HHT; Santa Cruz) treatments were performed at the indicated concentrations compared to 0.1% DMSO for 72 h or as indicated. Combination treatments of HHT with imatinib mesylate (in DMSO; LC Laboratories) or ripretinib (in DMSO; MedChem Express) were carried out at a fixed, equal ratio, as suggested by Chou and Talalay^35^. Single agent treatments with imatinib or sunitinib (in DMSO; LC Laboratories) served as positive control. A concentration of 1 µM was chosen to serve as a reference point for the ability to compare our results to previous studies that also used this concentration for their control treatments^15,18,19,21,36–40^. Cycloheximide (CHX) treatment was performed at 30 µg/ml in dH_2_O unless otherwise noted.

### Immunological and cell staining methods

Protein lysates of cells growing as monolayer were prepared as described previously^15^. In brief, cells were scraped into lysis buffer (1% NP-40, 50 mM Tris-HCl pH 8.0, 100 mM sodium fluoride, 30 mM sodium pyrophosphate, 2 mM sodium molybdate, 5 mM EDTA, 2 mM sodium orthovanadate) with protease inhibitors (10 µg/ml aprotinin, 10 µg/ml leupeptin, 1 µM phenylmethylsulfonyl fluoride). The suspension was incubated with shaking (1 h, 4°C) and cleared by centrifugation (13,000 rpm, 30 min, 4°C). The Bradford assay (Biorad) was used to determine protein concentrations, and 30 µg of protein were loaded on 4-12% Bis-Tris gels (Invitrogen).

Primary antibodies used for immunoblotting were p4E-BP1 T37/46, 4E-BP1 (both Cell Signaling), actin (Santa Cruz), cleaved caspase 3 (Cell Signaling), cyclin A (Novocastra), pKIT Y719 (Cell Signaling), KIT (DakoCytomation/Agilent), p27^Kip1^ (BD Biosciences Pharmingen), PARP (Invitrogen), pS6K T389, S6K (both Abcam).

Immunohistochemistry was performed on sections (4 μm) of formalin-fixed, paraffin-embedded (FFPE) mouse tumors from *in vivo* experiments. Slides were deparaffinized, underwent antigen retrieval and were incubated with primary antibodies (KIT, DakoCytomation/Agilent; Ki-67, GeneTex; cleaved caspase 3, Cell Signaling). The signal was detected using biotin-conjugated secondary antibodies (Abcam), POD-streptavidin (Roche), and 3’diaminobenzidine-tetrahydrochloride (DAB) chromogen (Abcam). For Ki-67 and cleaved caspase 3 stains, the number of positive cells was counted in 10 high power fields (HFP) per tumor at 400-fold magnification, with an average of 245 cells counted per HPF.

### Measurement of protein synthesis

To analyze protein synthesis activity, cellular incorporation of L-homopropargylglycine (HPG), a methionine analogue with alkyne moiety was detected by Click-iT reaction with azide-modified Alexa 488^41^. Cells treated with HHT, imatinib or sunitinib for 1h or 8h were labelled with 25 µM HPG (Invitrogen, C10102) for the last 30 min of the drug treatment in L-methionine-depleted medium containing 5% dialyzed fetal calf serum (FCS; Gibco). Harvested cells were washed with PBS, fixed in 10% buffered formalin (4% formaldehyde) for 5 min at room temperature and permeabilized for 30 min in PBS containing 0.1% saponine (Sigma) and 1% FBS (HyClone). The Click-iT reaction was performed for 30 min at room temperature by suspending the cells in Click-iT cell reaction buffer containing 1 mM azide-modified Alexa 488 and 2 mM CuSO4 (Invitrogen). Cells were washed twice in PBS with 1% FCS, and the cell suspension was subjected to flowcytometry analysis. The protein synthesis inhibitor cycloheximide (100 µg/ml) was used as a positive control in this assay. Cells without Click-iT reaction were used as a background control.

### *In vitro* apoptosis and proliferation assays

Apoptosis and cell viability studies were performed using the Caspase-Glo® and CellTiter-Glo® luminescence-based assays (Promega) as described previously^19^. Cells were cultured in 96-well flat-bottomed plates and incubated with HHT (or DMSO-only solvent control) for 48h (Caspase-Glo®) or 72h (CellTiter-Glo®). Luminescence was measured with a BioTek Synergy 2 Luminometer (BioTek) and normalized to the DMSO-only control. IC_50_s were calculated using the AAT Bioquest IC50 calculator (https://www.aatbio.com/tools/ic50-calculator).

### TUNEL assay

Apoptotic cells were detected using the terminal deoxynucleotidyl transferase-mediated dUTP nick end labeling (TUNEL assay; Roche Applied Sciences) according to the manufacturer’s recommendations.

### Reverse transcriptase (RT)-PCR

RNA extraction (Qiagen RNeasy Mini kit) and RT-PCR was performed as described previously^18^. Exon-overlapping, mRNA/cDNA-specific primers were used to amplify *KIT* (forward: 5’-TCATGGTCGGATCACAAAGA-3’, reverse: 5’-AGGGGCTGCTTCCTAAAGAG-3’; Operon) or *β-actin* (forward: 5’-CCAAGGCCAACCGCGAGAAGATGAC-3’, reverse: 5’-AGGGTACATGGTGGTGCCGCCAGAC-3’), with *β-actin* serving as reference gene and loading control.

### GIST xenograft model

One million GIST-T1 cells were bilaterally injected in the flank of twelve female adult athymic nude mice (BALB/c nude mice; Charles River Laboratories^42^. When tumors were approximately 60 mm^3^ in size, mice were randomized into groups of 4 animals each for each treatment regimen. In a combined dose-finding/efficacy experiment, homoharringtonine (2 mg/kg or 4 mg/kg in DMSO) or placebo (solvent control) was administered intraperitoneally (i.p.) twice per week for three weeks. Initial doses were chosen as reported in Kantarjian et al.^25^ and Alvandi et al.^28^. However, 4 mg/kg was toxic to all animals after one dose and mice treated at 2 mg/kg also showed signs of toxicity. The latter were switched to 1 mg/kg after the initial treatment, which they tolerated well. The placebo-treated mice showed no signs of toxicity. Tumor volume, weight and general health of the mice were recorded. After the mice were sacrificed, tumors were excised, formalin-fixed and paraffin-embedded (FFPE) for histopathological examination and IHC. Mice were sacrificed using CO_2_ and cervical dislocation. The animal experiment was approved by the IACUC of the University of Pittsburgh. All experiments were performed in accordance with the relevant guidelines and regulations and in with ARRIVE guidelines.

### Statistical analysis

Statistical significance was assessed using the 2-tailed Student’s t-test for independent samples. P values ≤0.05 were considered significant. The Chou–Talalay combination index (CI) method was used to evaluate synergism between HHT and imatinib or ripretinib, respectively^35^. CI values were calculated using CompuSyn (https://www.combosyn.com/) version 3.0.1 and plotted against the fraction affected (Fa). CI values of <1 were considered to be synergistic, whereas values of >1 indicate antagonism and a CI of 1 an additive effect.

## RESULTS

### Homoharringtonine inhibits nascent protein synthesis and abrogates KIT protein expression in GIST cells

Homoharringtonine (HHT) is a first-in-class mRNA translation elongation inhibitor that is FDA-approved for the treatment of TKI-resistant CML^22–25,28^. To determine whether HHT inhibits protein biosynthesis in GIST cells, nascent protein synthesis was measured by using a fluorescence-based assay that measures incorporation of the methionine analogue L-homopropargylglycine (HPG) via Click-iT technology and detection with azide-modified Alexa 488 (**Fig. 1A**). Imatinib (IM)-sensitive (GIST882, GIST-T1) and IM-resistant (GIST48, GIST430) human GIST cell lines were treated with HHT in comparison to the mRNA translation inhibitor cycloheximide and either IM (GIST882, GIST-T1) or sunitinib (SU; GIST430, GIST48), current standard medications for IM-naïve and IM-resistant GIST, respectively. HHT treatment at 1 h and 8 h led to a substantial decrease in protein translation activity that was comparable to cycloheximide in all cell lines. While IM and SU also resulted in a reduction of protein synthesis, changes were not as pronounced and more noticeable at the 8 h time point and are therefore likely a secondary effect due to inhibition of the mTOR signaling axis downstream of KIT^15^.

**Figure 1.**
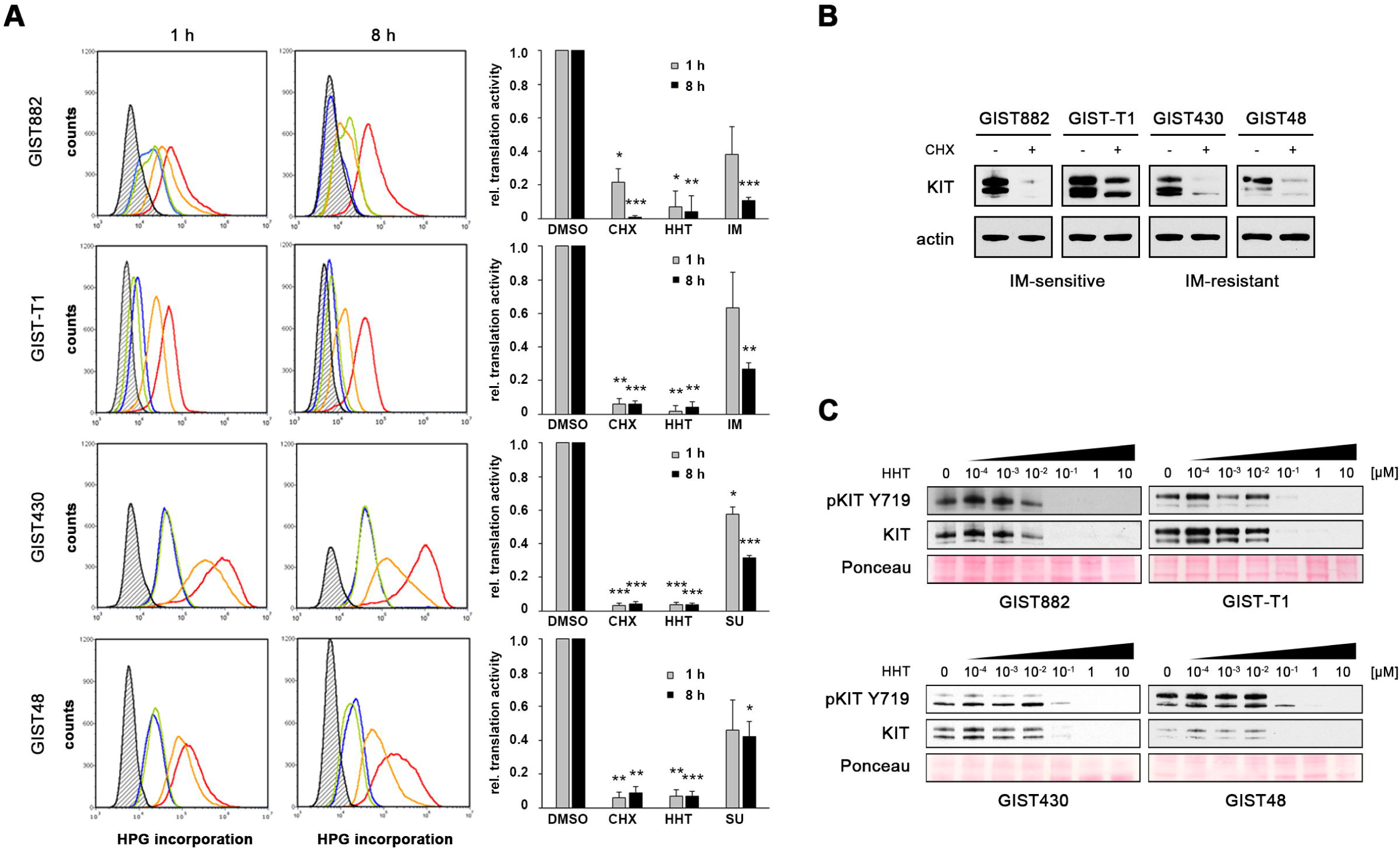
Homoharringtonine inhibits nascent protein synthesis and reduces KIT protein expression in GIST cells. **(A)** Nascent protein synthesis of GIST cells treated with homoharringtonine (HHT, 0.1 μM; green), cycloheximide (CHX, 100 μg/ml; blue), KIT inhibitor (imatinib [IM] 1μM for GIST882 and GIST-T1; sunitinib [SU] 1μM for GIST430 and GIST48; orange) or 0.1% DMSO control (red) for 1 h or 8 h. Cells were labelled with HPG during the last 30 min of drug treatment and incorporated cellular HPG linked to azide-modified Alexa488 was quantitated by flow cytometry. Unstained control cells are shown in black in the histogram. A change in nascent protein synthesis is indicated as a relative mean fluorescence value to the DMSO control in the bar graphs. Columns, mean + SE; *, p<0.05 in comparison to DMSO control; **, p<0.01 in comparison to control; ***, p<0.001 in comparison to control (Student’s t-test, 2-tailed). **(B)** Immunoblot analysis for KIT protein expression of IM-sensitive (GIST882, GIST-T1) and IM-resistant (GIST430, GIST48) GIST cells after treatment with the protein translation inhibitor cycloheximide (CHX; 30 μg/ml for 3 h). **(C)** Immunoblotting for KIT protein expression in IM-sensitive (GIST882, GIST-T1) and IM-resistant (GIST430, GIST48) GIST cells after treatment with HHT for 72 h at the indicated concentrations. **(B, C)** Grouped immunoblot images are either cropped from different parts of the same gel or from a separate gel run with another aliquot of the same protein lysate Abbreviations: 882 (GIST882), T1 (GIST-T1), 430 (GIST430), 48 (GIST48).

Many proteins derived from oncogenic driver genes have a high turnover rate and could hence be suitable targets for mRNA translation inhibitors such as HHT^22^. To determine whether this is the case for the mutant KIT protein in GIST cells, levels of KIT expression were determined after inhibition of protein translation with cycloheximide. KIT expression decreased to almost undetectable levels in most cell lines after 3 hours, indicating that the protein has a half-life of less than 1.5 hours in those cells (**Fig. 1B**). We then tested whether HHT has a similar effect and could show that treatment leads to a dose-dependent reduction of KIT protein expression in IM-sensitive (GIST882, GIST-T1) and IM-resistant (GIST430, GIST48) GIST cells when analyzed by immunoblotting (**Fig. 1C**).

### HHT induces apoptosis and cell cycle arrest in GIST cells

GIST cell survival is dependent on oncogenic KIT expression and activation^1,4,15–19^. To test whether the substantial downregulation of KIT protein expression after HHT treatment leads to a cytotoxic response, GIST cells were incubated with increasing concentrations of HHT over a 100,000-fold concentration range (0.0001 µM to 10 µM). Cell viability and apoptosis were measured using luminescence-based assays and showed robust changes in all cell lines tested (**Fig. 2A**). IC_50_s ranged from as low as 5 nM (GIST-T1) to 132 nM (GIST48; **Table 1**).

**Table 1.** IC_50_ (cell viability) values of homoharringtonine (HHT) in IM-sensitive (GIST882, GIST-T1) and IM-resistant (GIST430, GIST48) GIST cells.

|  | HHT IC<sub>50</sub> (nM) |
| --- | --- |
| **GIST882** | 32 |
| **GIST-T1** | 5 |
| **GIST430** | 107 |
| **GIST48** | 132 |

**Figure 2.**
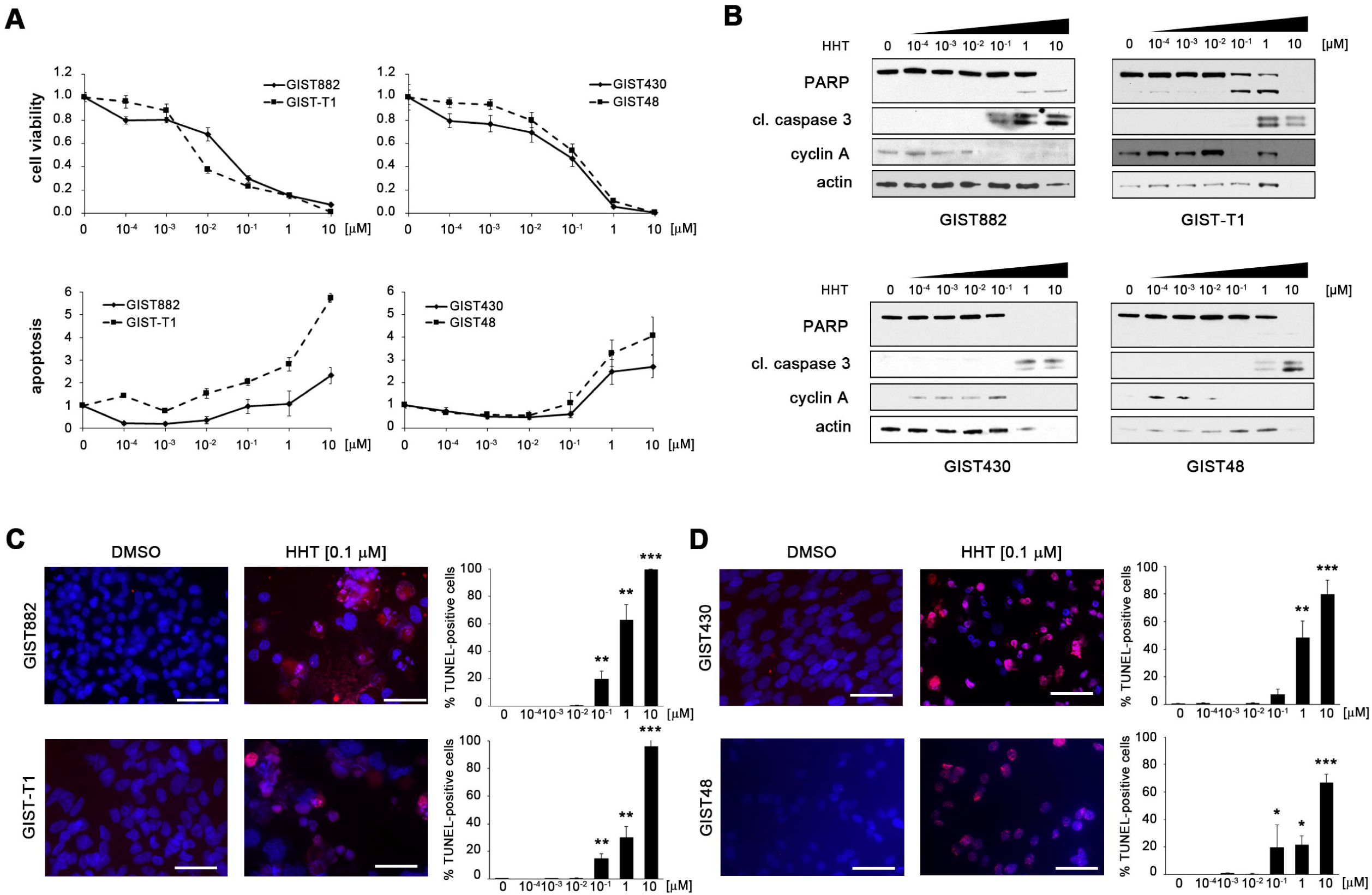
HHT has a pro-apoptotic effect in GIST cells and reduces cell viability in a dose-dependent manner. **(A)** Dose-dependent effect of HHT on cell viability (upper panels) and induction of apoptosis (lower panels) of IM-sensitive (GIST882, GIST-T1; left panels) and IM-resistant (GIST48, GIST430; right panels) GIST cells as measured by luminescence-based assays (mean□+/− □SE). **(B)** Immunoblot analysis for markers of apoptosis and cell cycle regulation after treatment of GIST cells with increasing concentrations of HHT (72 h) as indicated. Grouped immunoblot images are either cropped from different parts of the same gel or from a separate gel run with another aliquot of the same protein lysate. **(C, D)** Dose-dependent effect of HHT on induction of apoptosis in IM-sensitive **(C)** and IM-resistant **(D)** GIST cells as measured by TUNEL assay. Cells were treated for 72□h. Graphs represent mean and standard error of at least two experiments with an average of 500 cells counted per condition. Scale bar, 50 μm. Columns, mean + SE; *, p<0.05 in comparison to DMSO control; **, p<0.01 in comparison to control; ***, p<0.001 in comparison to control (Student’s t-test, 2-tailed).

As demonstrated by immunoblotting, HHT led to the induction of apoptosis (as shown by PARP and caspase 3 cleavage) and cell cycle exit (as shown by a reduction of cyclin A expression) at concentrations as low as 0.1 µM (**Fig. 2B**). The pro-apoptotic effect of HHT was further verified by terminal deoxynucleotidyl transferase-mediated dUTP nick end labeling (TUNEL) assay (**Fig. 2C, D**).

### The effect of HHT is time-dependent and leads to inhibition of translation initiation

Having shown that HHT induces apoptosis and cell cycle exit at a concentration as low as 0.1 µM, we used this concentration to determine the dynamic between translation inhibition and cellular response. Levels of total and phosphorylated (Y719) KIT rapidly decreased starting after 1 hour of treatment and were completely lost in all cell lines after 8 to 24 hours (**Fig. 3A**). Induction of apoptosis as indicated by PARP and caspase 3 cleavage as well as cell cycle exit (reduction of cyclin A levels) was delayed, starting at 24 hours after treatment in most cell lines (**Fig. 3B**).

**Figure 3.**
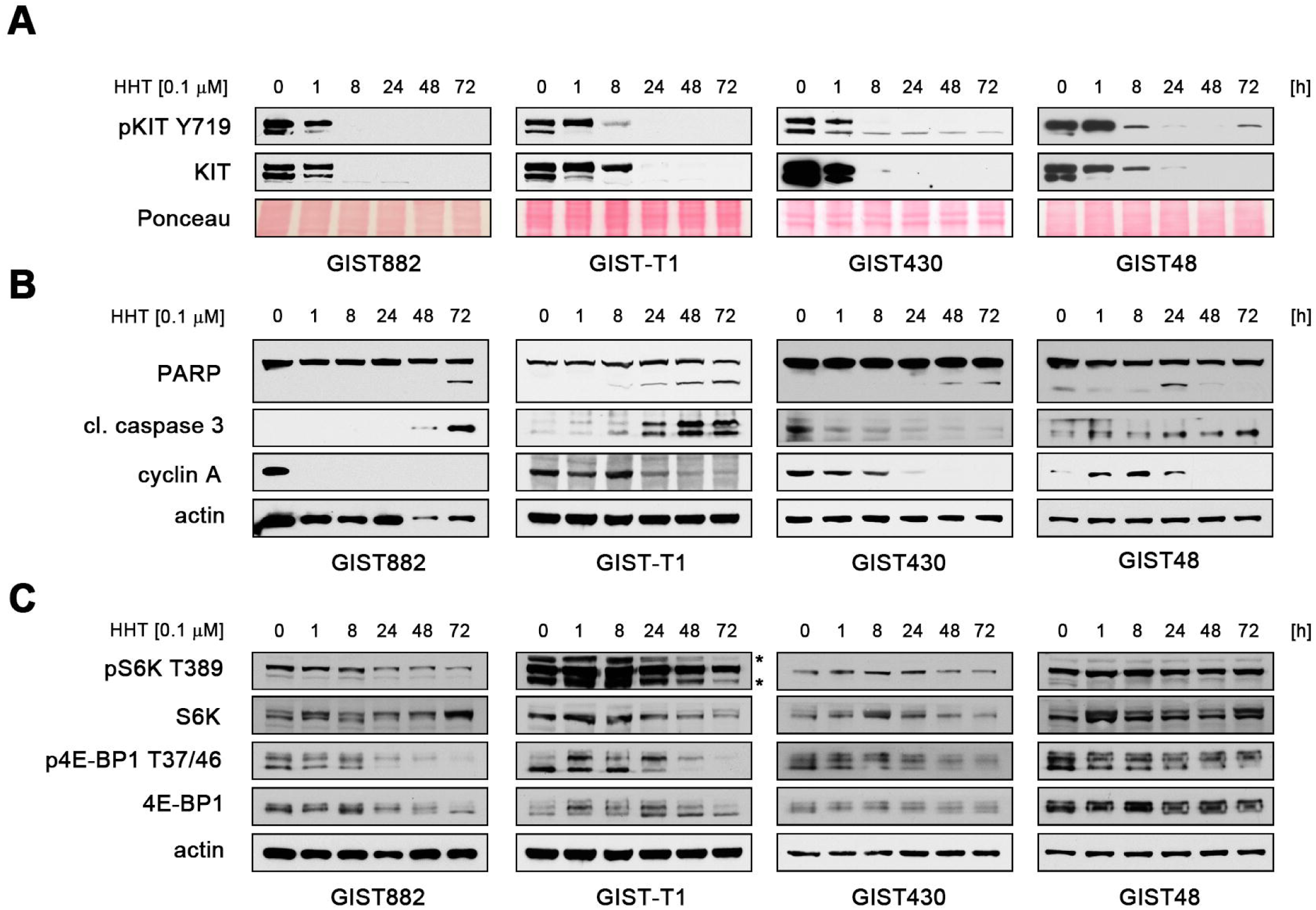
The effect of HHT is time-dependent and leads to secondary inhibition of translation initiation. **(A-C)** Immunoblot analysis of **(A)** KIT activation (phosphorylation at Y719) and protein expression, **(B)** markers of apoptosis and cell cycle regulation and **(C)** activation of the protein translation regulators ribosomal S6 kinase (S6K; T89) and 4E-BP1 (T37/46) in GIST cells after treatment with DMSO or HHT (0.1□μM) for the indicated times. * indicates unspecific band. Grouped immunoblot images are either cropped from different parts of the same gel or from a separate gel run with another aliquot of the same protein lysate.

HHT-treated cells also showed decreased expression and phosphorylation of ribosomal S6 kinase and 4E-BP1, a major regulator of translation initiation factor eIF4E (**Fig. 3C**). Both are direct targets of the mTOR kinase, which is downstream of KIT^15^. Taking into account that inhibition of KIT signaling leads to downregulation of nascent protein synthesis (**Fig. 1A**) and the fact that HHT interferes with the elongation step of protein translation, the above findings likely represent a secondary effect after loss of KIT expression/activation and are not a direct effect of HHT on the translational machinery.

### Loss of KIT expression after HHT treatment is not due to reduced mRNA transcription

To further prove that reduced KIT protein expression in GIST cells after HHT treatment is due to the inhibition of protein translation, we analyzed the effect of the drug on mRNA transcription. *KIT* mRNA levels did not or only minimally decrease with increasing duration of treatment (**Fig. 4**), thus further corroborating the notion that HHT’s main mechanism of action is the inhibition of protein translation.

**Figure 4.**
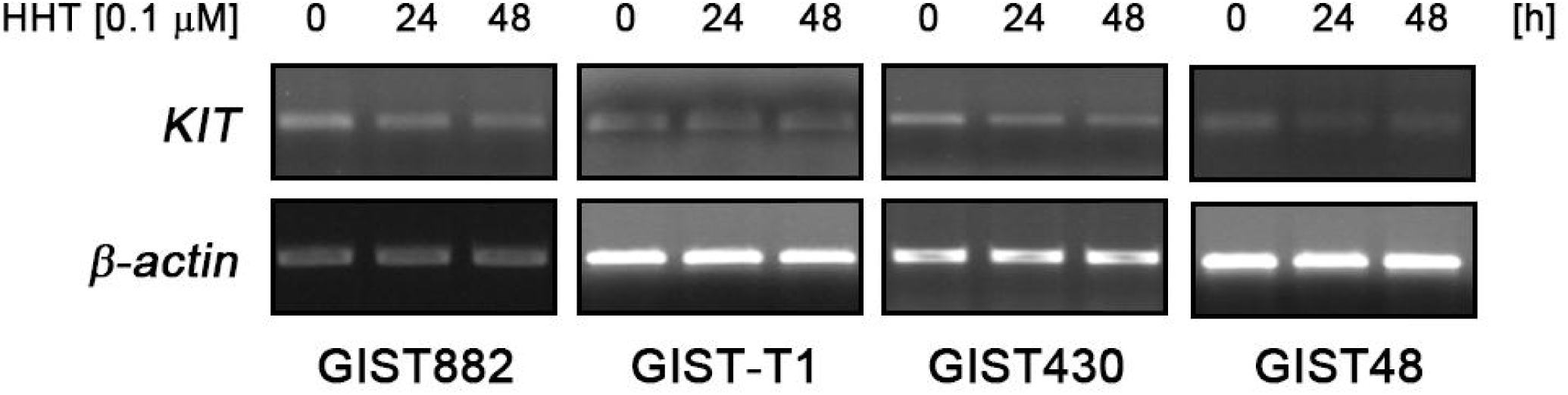
*KIT* mRNA levels are not affected by HHT treatment. RT-PCR of *KIT* mRNA expression after treatment of GIST cells with DMSO or 0.1 μM HHT for the indicated times. *β-actin* mRNA expression serves as loading control.

### HHT has *in vivo* antitumor activity in a GIST xenograft

To determine the antitumoral activity of HHT *in vivo*, we used a GIST xenograft model (GIST-T1). Combined dose finding and efficacy testing showed toxicity at higher doses. However, animals treated with low-dose HHT (1 mg/kg), showed a remarkable tumor shrinkage. This dose is approximately equivalent to half of the standard human dose, which was shown to result in a maximal plasma concentration (C_max_) of 36.2 ng/ml (equivalent to ∼120 nM)^43,44^. Over the course of three weeks, HHT treatment resulted in a marked histopathologic response with large areas of central necrosis (**Fig. 5A**, upper panels). As expected from our *in vitro* studies, HHT-treated tumor cells showed a loss of KIT expression (**Fig. 5A**, lower panels). This was accompanied by significant tumor reduction, while placebo-treated xenografts continued to grow exponentially (**Fig. 5B**). Of note, the response to IM treatment is characterized by disease stabilization in most cell line-derived GIST xenograft models^45–47^, indicating that HHT has a considerably higher cytotoxic activity. Furthermore, HHT led to a significant reduction of proliferation (**Fig. 5C**; reduction of the Ki-67-positive proliferation fraction from 36.4% in placebo-treated controls to 12.8% in HHT-treated tumors) and a significantly increased proportion of apoptotic cells (cleaved caspase 3-positive) from 6.3% in vehicle-treated controls to 14.6% in HHT-treated animals (**Fig. 5D**). Taken together, HHT has significant activity in GIST xenografts.

**Figure 5.**
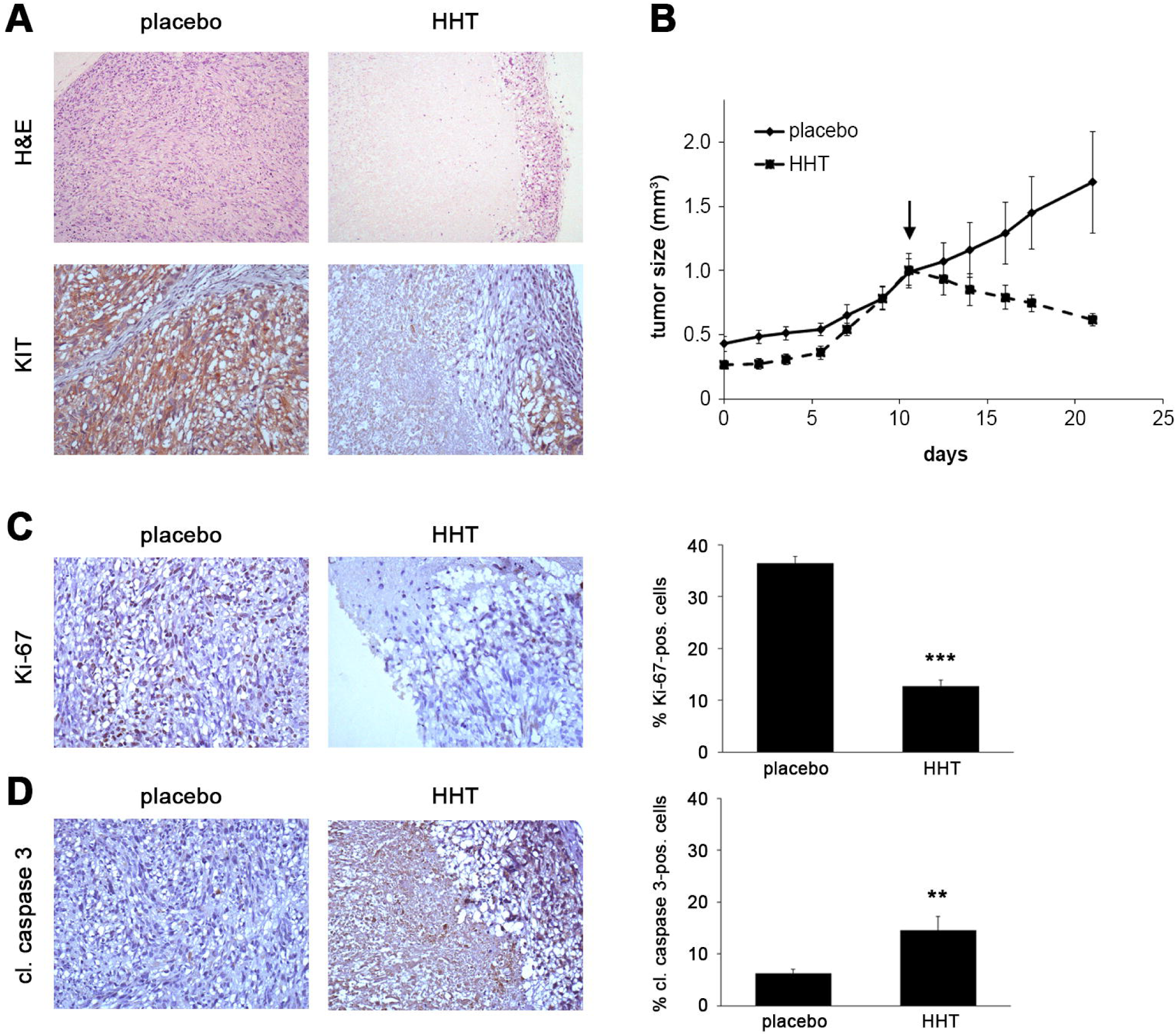
HHT is effective in a GIST xenograft. **(A)** Histopathologic response (hematoxylin and eosin, H&E; 10X; upper panels) and immunohistochemical staining for KIT (20X; lower panels) of GIST-T1 xenografts after treatment with HHT in comparison to placebo-treated controls. **(B)** Relative tumor volume of GIST-T1 xenografts since inoculation and over the course of a 21-day treatment with vehicle (solid line) or HHT (dashed line). Measurements represent the average of four tumors per group treated either with placebo or 2 mg/kg HHT (reduced to 1 mg/kg at day 7) for 21 days. Arrow indicates start of treatment. **(C**,**D)** Immunohistochemical staining for **(C)** Ki-67 (20X; upper panels) and **(D)** cleaved caspase 3 (20X; lower panels) of GIST-T1 xenografts treated with HHT or placebo control. Graphs represent mean and standard error of positive cells in 10 high power fields (HPF; 400-fold magnification), with an average of 245 cells counted per HPF. Columns, mean + SE; **, p□≤ □0.01 in comparison to control; ***, p□≤ □0.001 in comparison to placebo control (Student’s t-test, 2-tailed).

In summary, we show that HHT is effective in IM-sensitive and IM-resistant GIST cells as well as a GIST xenograft model. Its mechanism of action involves a substantial downregulation of KIT expression and subsequent inhibition of downstream signaling pathways and induction of apoptosis through inhibition of nascent protein biosynthesis.

## DISCUSSION

GISTs can be successfully treated with tyrosine kinase inhibitors (TKIs) that target KIT, such as imatinib mesylate, but many patients develop resistance to imatinib as well as to second- or later-line TKIs^5,6^. Mechanistically, TKI resistance in GIST is largely mediated through secondary *KIT* mutations, indicating that active KIT continues to be the main driver of the disease during tumor progression^7,8^. Because of the intra- and intertumoral heterogeneity of those mutations, treatment responses may be uneven and lead to incomplete responses^20^. Targeting KIT by inhibiting protein translation is independent of the *KIT* mutational status and may hence lead to more complete and durable therapeutic effects.

In this study, we show that homoharringtonine (HHT), a first-in-class inhibitor of protein biosynthesis at the step of peptide chain elongation, has significant antitumoral activity in GIST cells. This was independent of the *KIT* mutational status and imatinib sensitivity. Evaluating several human GIST cell line models, HHT potently inhibited nascent protein synthesis, KIT protein expression and phosphorylation as well as downstream signaling. Apoptosis was induced in a dose- and time-dependent manner via induction of PARP and caspase 3/7 cleavage. At the same time, decreased levels of cyclin A point to HHT-induced cell cycle arrest in S phase. As expected, expression of *KIT* mRNA expression was largely unaffected. Results from a GIST xenograft model show antitumor activity of HHT *in vivo*. Taken together, our results indicate that HHT has potent antitumor activity in GIST *in vitro* and *in vivo* and encourage clinical studies in GIST patients.

The fact that HHT has activity in imatinib-resistant cells makes it a promising medication for use in the TKI-resistant setting. Although two new compounds have been recently approved (ripretinib [Qinlock®; Deciphera] and avapritinib [Ayvakit®; Blueprint Medicines]), resistance to both drugs still occurs^12–14,37^, and further therapeutic options are needed. In addition, avapritinib’s indication only covers PDGFRA D842V-mutant GIST, limiting its use to a very small patient population. Based on its mechanism of action, it is conceivable that HHT is also effective in *PDGFRA*-mutant GIST^3^, including mutations other than D842V, for which only few therapeutic options are available. Indeed, other cancers that are driven by oncogenically activated kinases, such as EGFR- or ALK-mutated non-small cell lung cancer and BRAF V600E-mutated melanoma should also be amenable to this approach^48^.

Mechanistically, it is highly likely that the loss of KIT expression induced by HHT is the main mediator of GIST cell death. GIST cells are dependent on active KIT signaling for their survival, and we have previously shown that knocking down KIT expression with small interfering RNA (siRNA) leads to a pronounced apoptotic response via a caspase-3-dependent mechanism^16,38^. Similarly, Jin et al. reported significantly increased apoptosis after silencing KIT in two *KIT*-mutant mastocytosis models^31^. Likewise, these cells were highly sensitive to treatment with HHT. In our study, loss of KIT expression induced by HHT also led to secondary changes that occurred delayed when compared to loss of KIT expression, but likely contributed to drug efficacy. For example, decreased phosphorylation of ribosomal S6 kinase and the translation initiation factor 4E-BP1, both downstream of the KIT/mTOR signaling axis, point to an inhibition of translation initiation in addition to HHT-mediated inhibition of the elongation step of protein translation^15^. Other factors of HHT-mediated apoptosis, such as signal transducer and activator of transcription (STAT) proteins, have been reported ^49,50^. However, the respective STAT proteins are either not expressed or not regulated in a KIT-dependent fashion in the models used in our current study^15^.

Data from the RTOG S0132 trial demonstrated that neoadjuvant therapy is associated with decreased KIT expression^51,52^, and this phenomenon has also been described in a subset of imatinib-resistant, progressing GISTs that were surgically removed^53–55^. Based on the results presented here, it would not be expected that these “KIT-negative” GISTs are sensitive to HHT. However, there is a possibility that KIT-negative GIST cell populations emerging under selection pressure over time rely on other oncogenic driver events for their survival which keep them amenable to HHT treatment. Indeed, we could show that GIST48B, a spontaneously derived subline of GIST48 that does not express KIT transcript and protein at detectable levels, is not entirely resistant to HHT (Suppl. Fig. 1). These results indicate that HHT has KIT-independent mechanisms in these cells. In contrast, it can be speculated that inherently KIT^low^ subpopulations, such as those that have been identified in mouse and human GIST models^52,56^, are likely not sensitive to HHT beyond a general sensitivity to global inhibition of protein synthesis.

Clinical trials to evaluate HHT’s efficacy in GIST patients are warranted and feasible. The drug is FDA-approved for TKI-resistant CML, and IC_50_ values of preclinical GIST models (55 – 132 nM) are comparable to those reported of CML *in vitro* models (20 – 150 ng/ml, equivalent to 67 – 500 nM)^57,58^. Current HHT dosing schedules include subcutaneous self-application as opposed to intravenous infusion^26^, which might be of importance to a patient population that is accustomed to oral anti-cancer medications. Adverse reactions, such as pancytopenia and diarrhea, have been reported in CML and might be due to HHT’s mechanism of action as a global inhibitor of protein biosynthesis^59^. One possibility to reduce these undesirable effects could be to combine HHT treatment with approved TKIs, thus potentially allowing for a dose reduction of each drug. In fact, preliminary results from our laboratory show synergism between HHT and imatinib (in IM-sensitive GIST cells) as well as ripretinib (in IM-resistant cells; Suppl. Fig. 2). It is still possible, however, that preclinical results do not translate into the clinic. Clinically solid tumors, such as GIST, sometimes respond differently to the same drug compared to hematological malignancies, as has been reported for the proteasome inhibitor bortezomib (Velcade®)^60,61^.

Taken together, we have shown that the protein translation inhibitor HHT is highly effective in GIST cells *in vitro* and *in vivo* and leads to a loss of KIT expression. Further studies to bring the drug into the clinic are warranted.

## DATA AVAILABILITY

All data generated or analyzed during this study are included in this published article and its Supplementary Information files.

## ACKNOWLEDGEMENTS

This work was supported by the GIST Cancer Research Fund (to A.D.), The Life Raft Group (A.D.), the Out of the Woods Foundation (A.D.), Pittsburgh Cure Sarcoma (A.D.), NIH P50CA121973 (University of Pittsburgh Skin Cancer Specialized Program of Research Excellence; M.S.), the Hillman Foundation (M.S.), a Klionsky Fellowship (Department of Pathology, University of Pittsburgh School of Medicine; to C.L.) and private donations (A.D.). This project used the UPMC Hillman Cancer Center and Tissue and Research Pathology/Pitt Biospecimen Core shared resource which is supported in part by award P30CA047904. A.D. and M.S. are supported by UPMC Hillman Cancer Center and in part by a grant from the Pennsylvania Department of Health. The Department specifically disclaims responsibility for any analyses, interpretations or conclusions. The authors would like to thank Jonathan A. Fletcher for sharing important reagents, Shou-Jiang Gao for translation of an article not available in English, Stefan Duensing for critically reading the manuscript and members of the Pittsburgh Sarcoma Research Collaboration (PSaRC) for invaluable discussions and support.

## AUTHOR CONTRIBUTIONS

Conception and design: A.D.; Development of methodology: D.M.L., S.P., A.R., L.L., P.T., A.A.A., C.L., J.L.R., F.S., M.S., A.D.; Acquisition of data: D.M.L., A.S., S.P., A.R., L.L., P.T., A.A.A., C.L., J.L.R., A.K., F.S., N.A., M.S., A.D.; Analysis and interpretation of data: D.M.L., A.S., S.P., A.R., P.T., A.A.A., J.L.R., C.L., F.S., N.A., M.S., A.D.; Writing, review and/or revision of the manuscript: D.M.L., A.S., S.P., L.L., A.R., C.L., P.T., A.A.A., C.L., J.L.R., L.P., N.A., M.S., A.D.; Administrative, technical, or material support: A.S., L.P., A.D.; Study supervision: A.D.

