## Supplemental Data for "Targeting the translational machinery in gastrointestinal stromal tumors (GIST) – a new therapeutic vulnerability"

### SUPPLEMENTARY FIGURE LEGENDS

**Supplementary Figure S1. HHT has activity in KIT-negative GIST48B cells.**

**(A)** RT-PCR of *KIT* mRNA expression of GIST48B cells in comparison to GIST48 after treatment with DMSO or 0.1 uM HHT for the indicated times.

**(B)** Immunoblot analysis for KIT protein expression of GIST48B cells in comparison to GIST48 after treatment with the protein translation inhibitor cycloheximide (CHX; 30 ug/ml for 3 h). Asterisk denotes an unspecific band.

**(C)** Nascent protein synthesis of GIST48B cells treated with homoharringtonine (HHT, 0.1 µM; green), cycloheximide (CHX, 100 µg/ml; blue), the KIT inhibitor sunitinib (SU, 1µM; orange) or 0.1% DMSO control (red) for 1 h or 8 h. Cells were labelled with HPG during the last 30 min of drug treatment and incorporated cellular HPG linked to azide-modified Alexa488 was quantitated by flow cytometry. Unstained control cells are shown in black in the histogram. A change in nascent protein synthesis is indicated as a relative mean fluorescence value to the DMSO control in the bar graphs. Columns, mean + SE; **, $p \leq 0.01$ in comparison to control; ***, $p \leq 0.001$ in comparison to DMSO control (Student's t-test, 2-tailed).

**(D)** Dose-dependent effect of HHT on cell viability (left panel) and induction of apoptosis (right panel) of GIST48B cells as measured by luminescence-based assays (mean +/− SE).

**(E)** Immunoblot analysis for markers of apoptosis and cell cycle regulation after treatment of GIST48B cells with increasing concentrations of HHT (72 h) as indicated.

**(F)** Immunoblot analysis for markers of apoptosis and cell cycle regulation after treatment of GIST48B cells with DMSO or HHT (0.1 µM) for the indicated times.

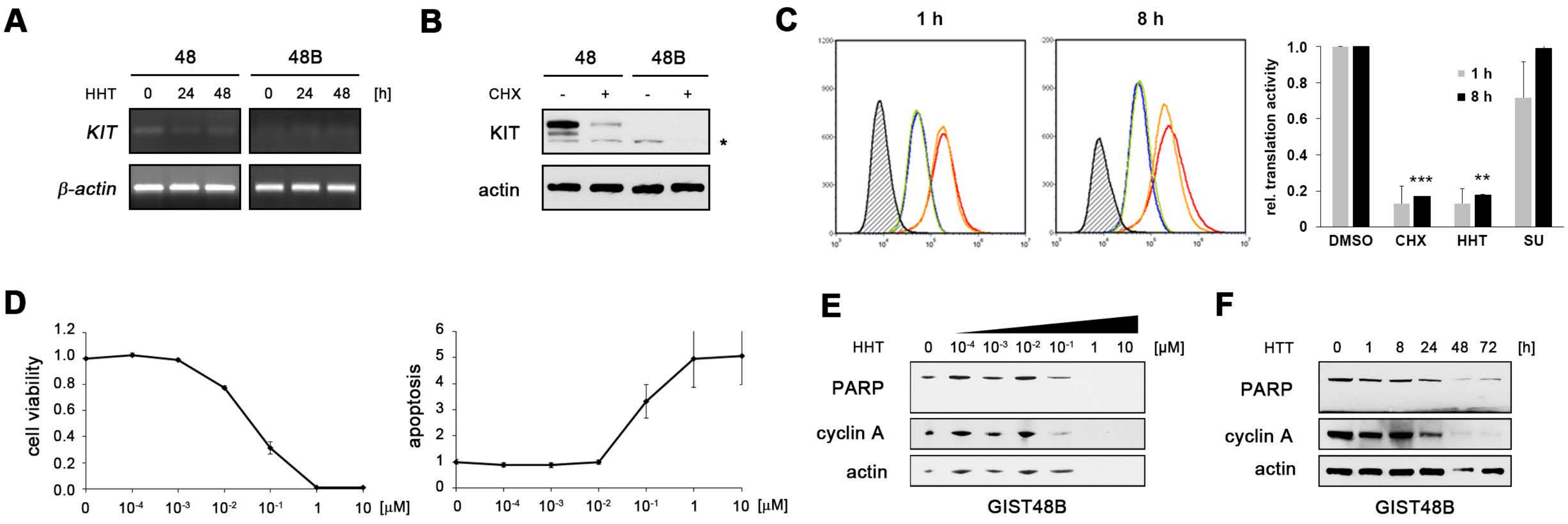


Lee *et al.*, Supplemental Figure 1

**Supplementary Figure 2. Combining HHT treatment with FDA-approved TKIs has a synergistic effect.**

Dose-dependent effect of HHT in combination with imatinib (IM; GIST882 and GIST-T1) or ripretinib (RIP; GIST430) on cell viability of IM-sensitive (GIST882, GIST-T1) and IM-resistant (GIST430) GIST cells as measured by luminescence-based assays (panels on the left; mean +/− SE). The combination index (CI; panels on the right) was calculated according to the method of Chou and Talalay [35] and plotted according to the fraction of cells affected. CI values of <1 indicate synergism of the two drugs, whereas values of >1 indicate antagonism and a CI of 1 an additive effect.

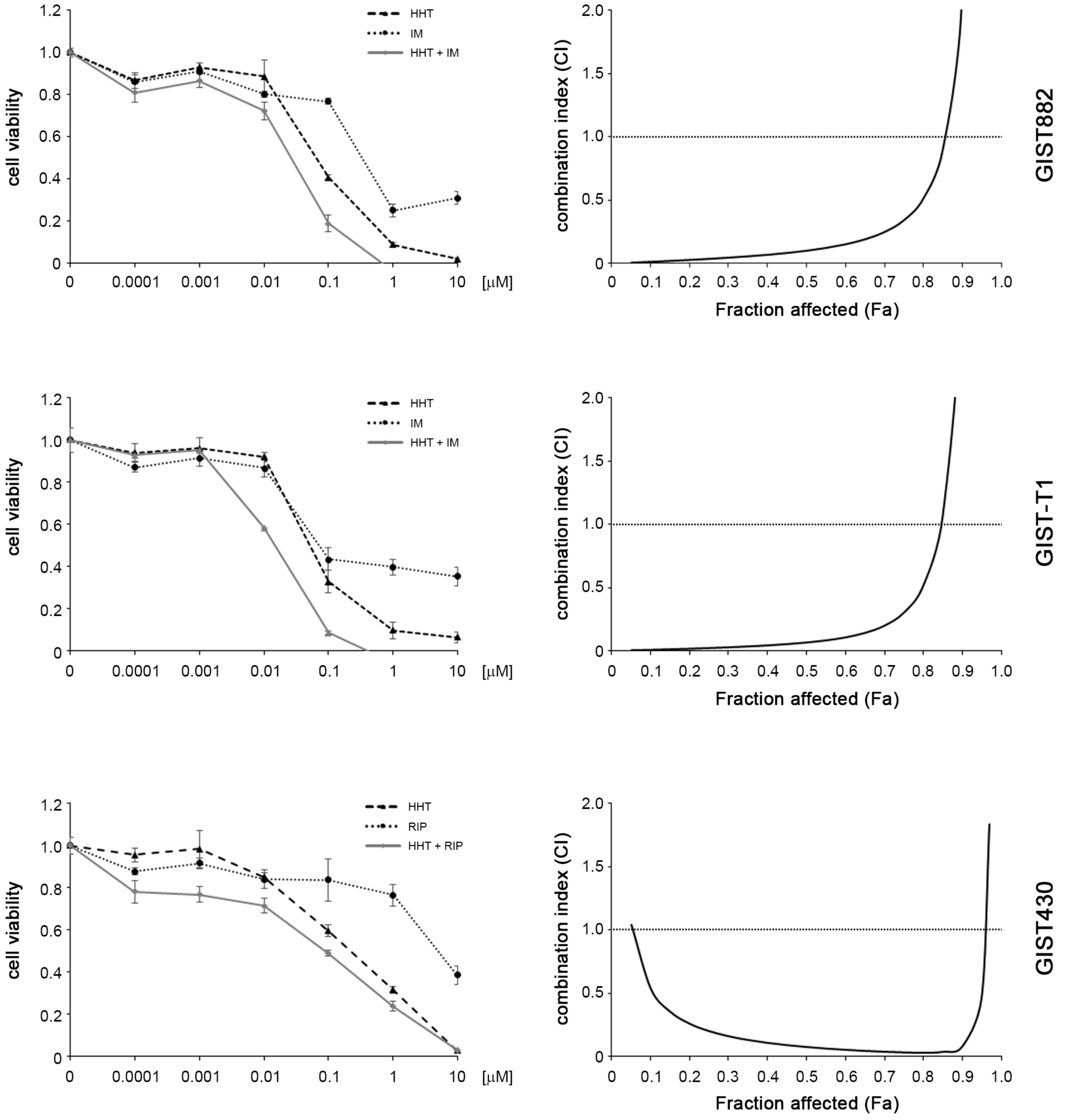


Lee et al., Supplemental Figure 2

**Supplementary Figure S3. Compilation of full blot images.**

Each page corresponds to a sub-figure of the manuscript (as noted in the lower right corner of the page) that contains a cropped blot image. On the respective page, either the entire blot or the entire region of the membrane that was stained for the respective marker is shown. Lanes, treatments and stains are labeled as in the manuscript figures, size markers are labeled in kDa.

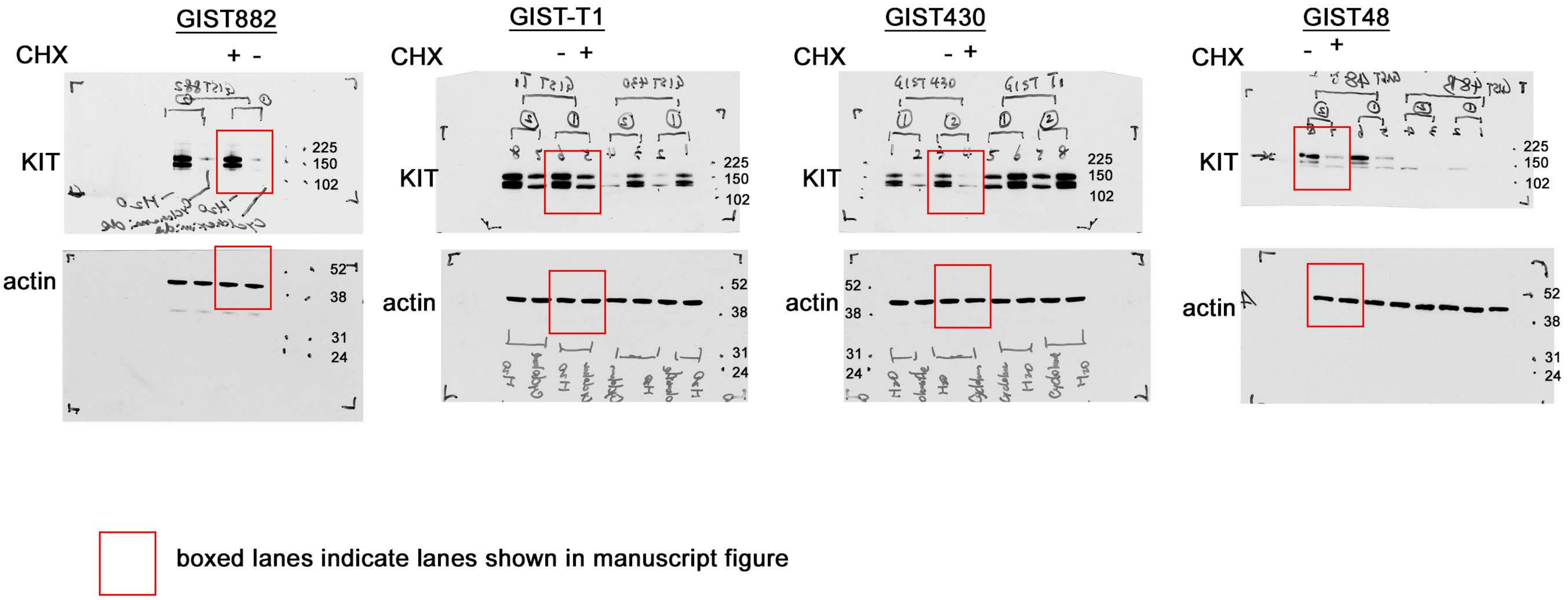


boxed lanes indicate lanes shown in manuscript figure

full blot images - Lee *et al.*, Figure 1B

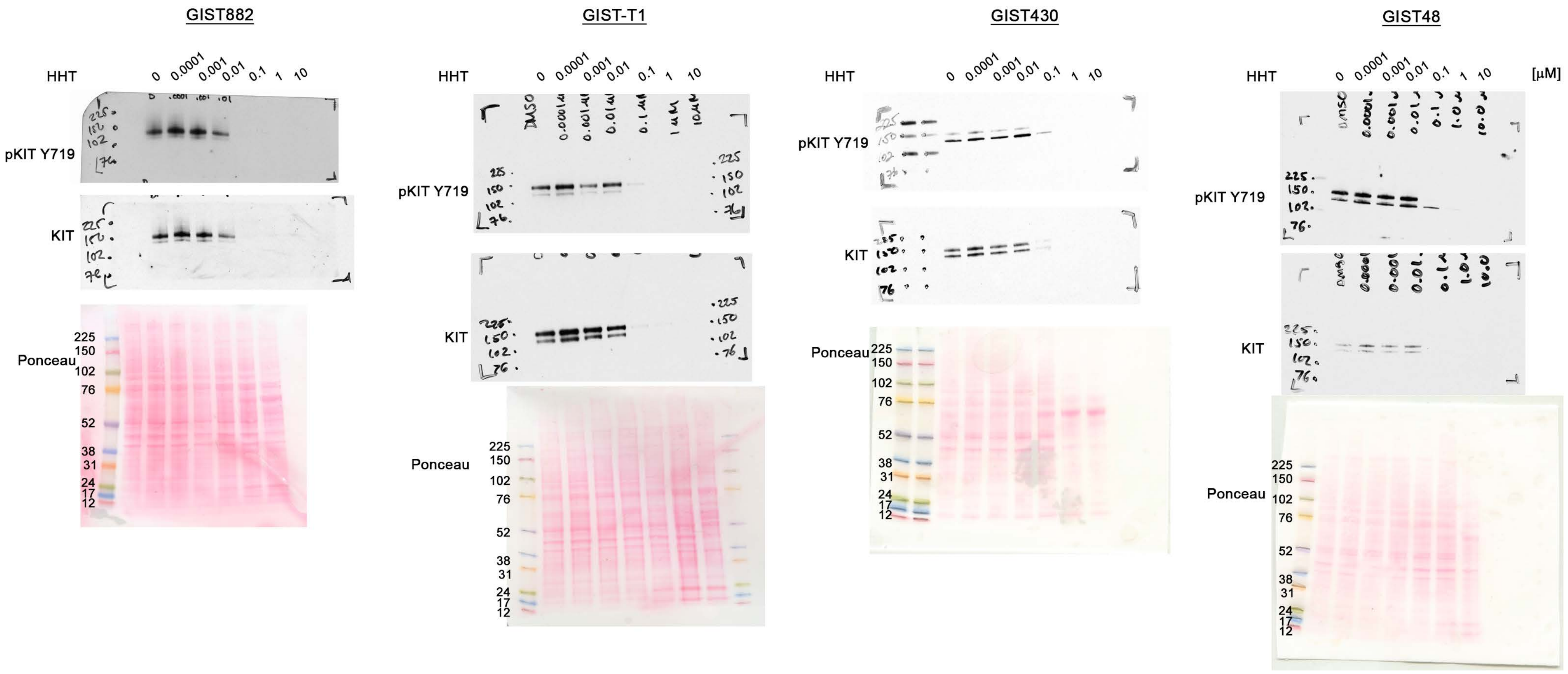


full blot images - Lee *et al.*, Figure 1C

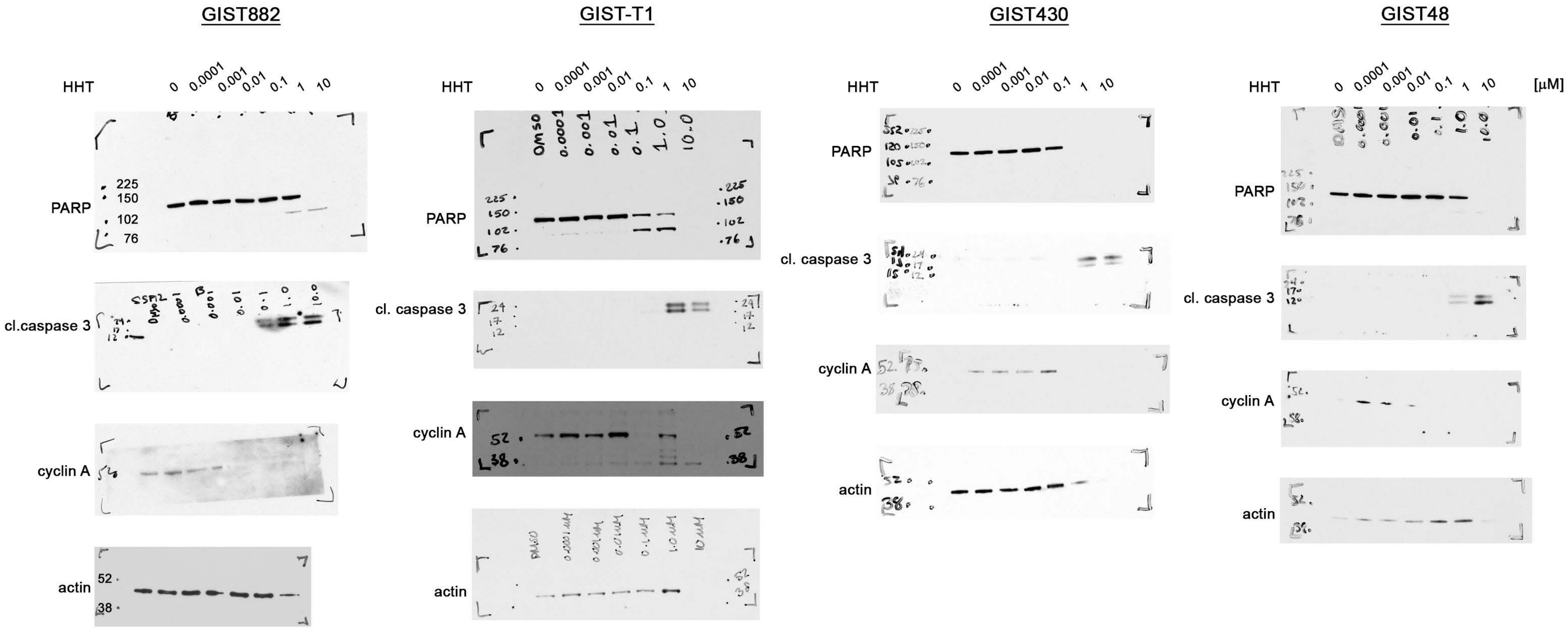


full blot images - Lee *et al.*, Figure 2B

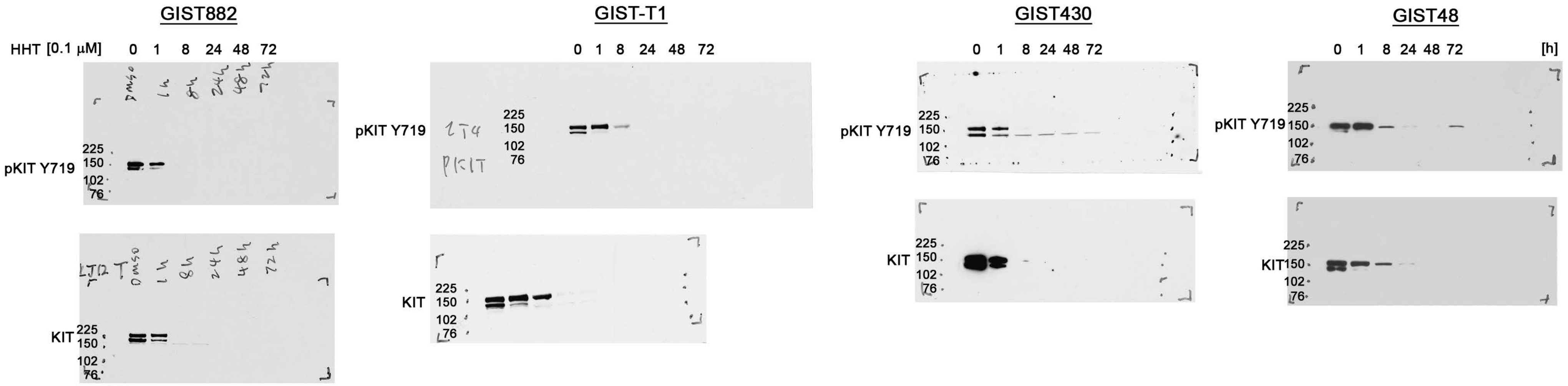


full blot images - Lee *et al*., Figure 3A

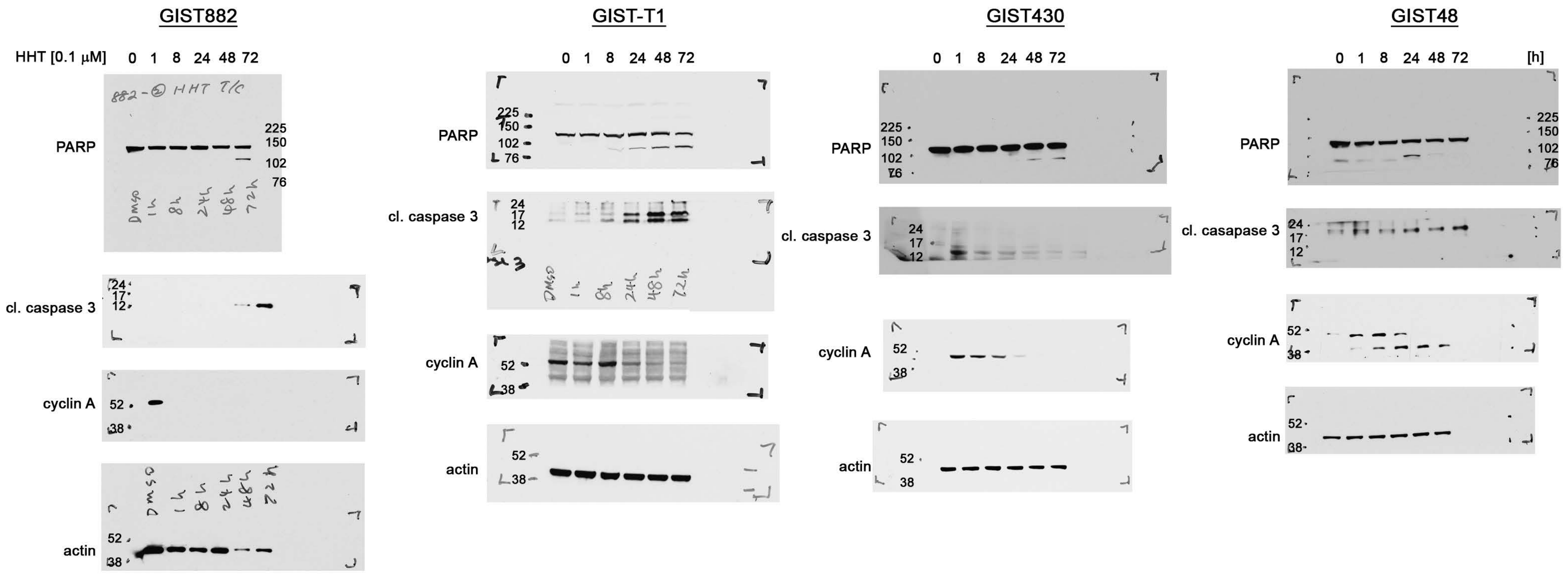


full blot images - Lee *et al.*, Figure 3B

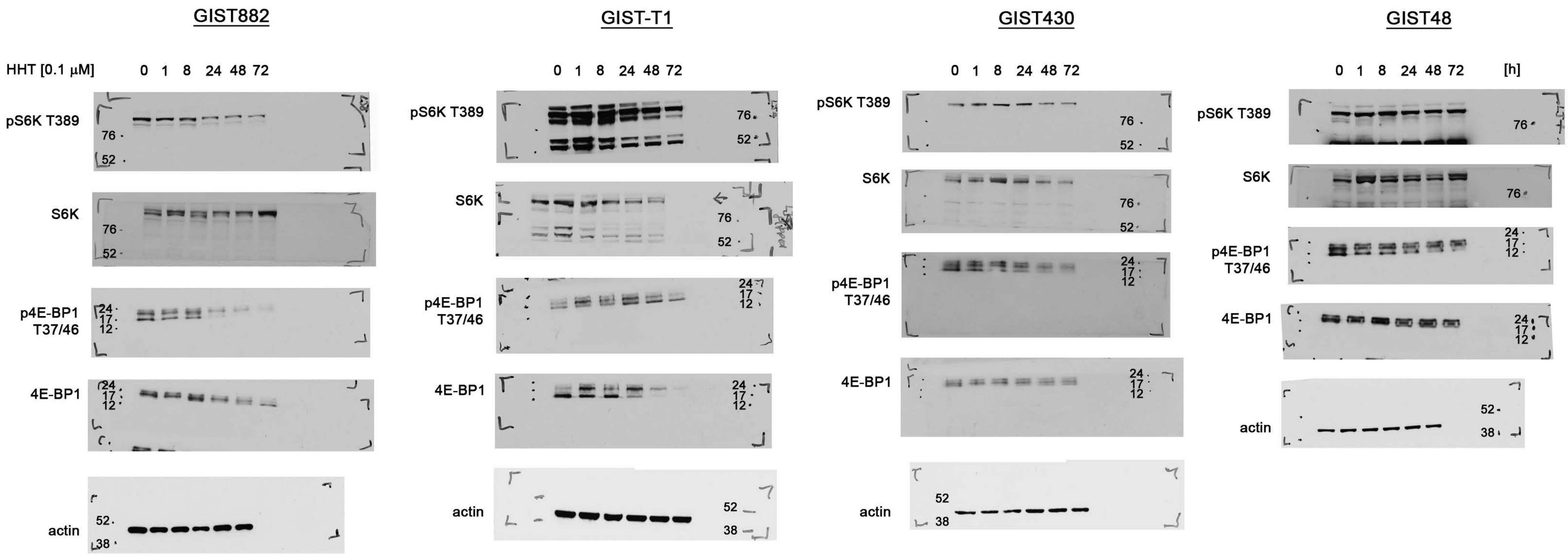


full blot images - Lee *et al.*, Figure 3C

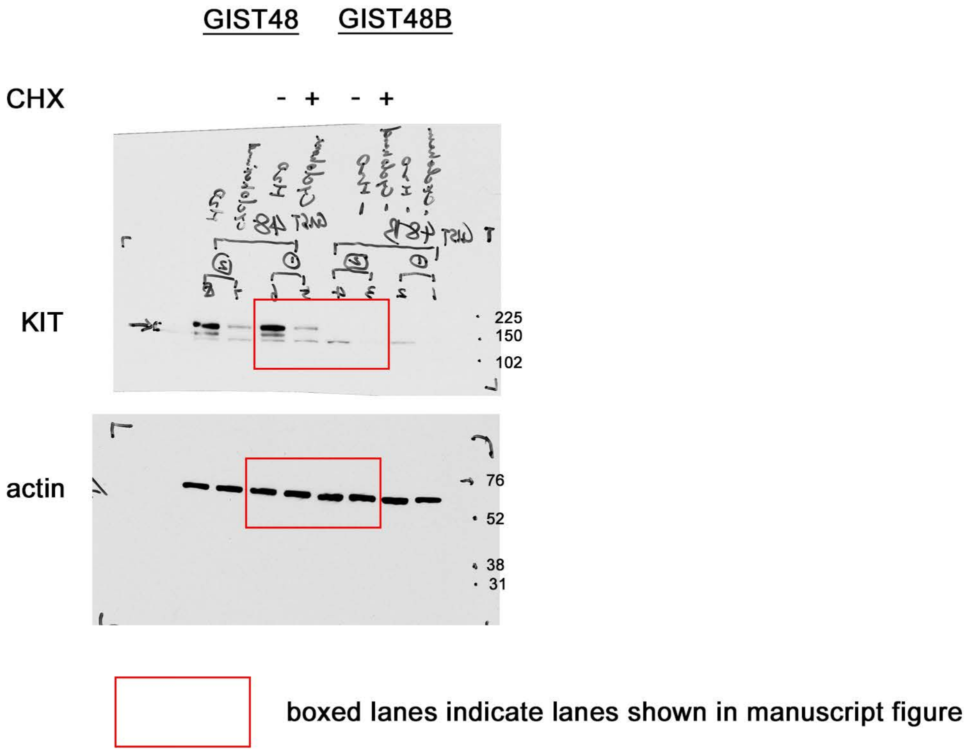


full blot images - Lee *et al.*, Supplemental Figure 1B

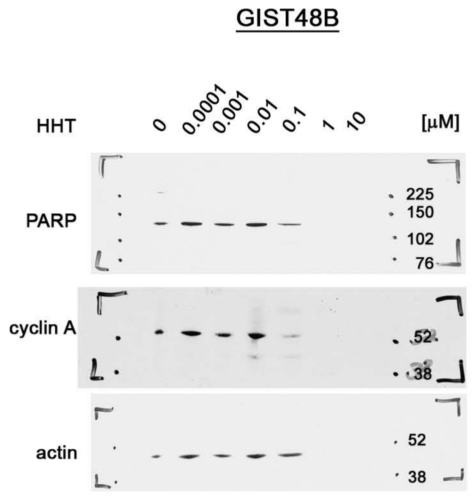


full blot images - Lee *et al.,* Supplemental Figure 1E

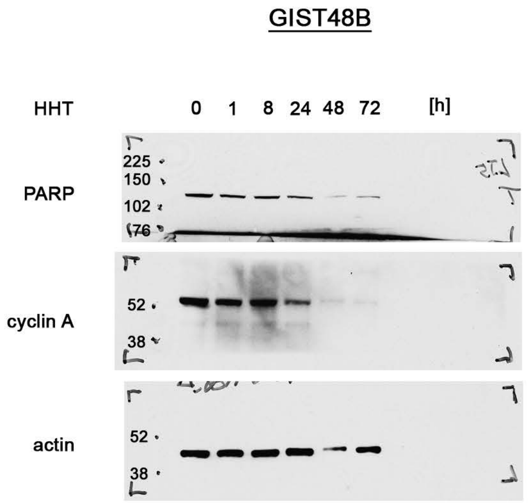


full blot images - Lee *et al.*, Supplemental Figure 1F
